# Evolution of the reproductive interactome: proteomic divergence and the origins of postmating-prezygotic reproductive incompatibilities

**DOI:** 10.1101/2025.07.14.664647

**Authors:** Jeremy M. Bono, Luciano M. Matzkin, Helen K. Pigage, Toan Hoang, Santana L. Navarrette, Marissa K. Benavidez, Clinton Green, Carson W. Allan

## Abstract

Reproductive proteins evolve rapidly, making them strong candidates for driving postmating-prezygotic (PMPZ) reproductive incompatibilities between populations. While most previous studies have focused on protein sequence divergence as a likely driver of PMPZ incompatibilities, reproductive proteomes may also diverge compositionally and/or quantitatively. Here, we combine quantitative proteomics, molecular evolutionary analyses, and protein-protein interaction (PPI) modeling to predict the molecular basis of reproductive incompatibilities between *Drosophila mojavensis* and *D. arizonae*. We demonstrate multidimensional divergence in reproductive proteomes including changes at the sequence, compositional, and quantitative levels. We further demonstrate that three divergent male seminal fluid proteins affect the size of the insemination reaction mass and/or fertilization success in *D. arizonae*. Despite high sequence divergence, predicted protein-protein interactions involving a conserved set of proteases and/or protease inhibitors were predicted to be maintained in heterospecific crosses. In contrast, predicted interspecies protein incompatibilities arose from proteome compositional divergence, suggesting that such changes may play a disproportionate role in PMPZ incompatibilities, at least for the subset of the interactome that we tested. Furthermore, extensive quantitative divergence, particularly for proteases and inhibitors, suggests pervasive stoichiometric mismatches in heterospecific matings. Altogether, our findings indicate that reproductive proteins are evolutionarily labile at multiple levels, and that compositional and quantitative divergence, rather than sequence changes alone, may be central to the early evolution of reproductive isolation.

## Introduction

Postmating molecular interactions between the sexes drive complex behavioral and physiological responses in females that determine reproductive outcomes [1]. Given their direct role in mediating successful reproduction, divergence in postmating molecular interactions between populations is especially likely to contribute to the speciation process [2]. In fact, postmating-prezygotic (PMPZ) reproductive isolation has been increasingly recognized as a common, rapidly evolving barrier to successful reproduction between diverging groups [2–5]. Despite this, progress in identifying the molecular basis and evolutionary forces driving PMPZ incompatibilities has trailed behind the study of other types of reproductive isolation [2].

The widespread observation that reproductive proteins evolve rapidly has led to the hypothesis that pervasive postcopulatory sexual selection and sexual conflict drives rapid divergence in reproductive proteins [5–8], and thus likely play crucial roles in generating PMPZ incompatibilities [2,3,5]. Most previous comparative studies of male and female reproductive proteomes have focused on protein sequence divergence, with the assumption that PMPZ incompatibilities arise from the disruption of coevolved molecular interactions through lineage-specific changes in protein sequence [6–9] (Fig. 1; scenario 1). However, some studies have also suggested that most sequence divergence in reproductive proteins is non-adaptive [10–12]. Moreover, relatively little attention has been directed other forms of reproductive proteome divergence that might also contribute to PMPZ isolation. For example, the male-female reproductive interactome could diverge through gains or losses of interactions that result from compositional changes of reproductive proteomes (i.e. presence/absence of specific proteins) (Fig 1; scenario 2A). In fact, *Drosophila* lineage-specific genes that arise *de novo* or through gene duplication are known to play fundamentally important roles in reproductive processes [13–20]. The male-female interactome could also diverge if conserved reproductive proteins gain or lose interacting partners in different lineages (Fig 1; scenario 2B). In either case, lineage-specific changes to the interactome may cause disruption of the normal postmating response in interpopulation crosses. Finally, reproductive proteins could also diverge quantitatively, resulting in stoichiometric mismatches between males and females from divergent populations (Fig 1; scenario 3). Although interactome and/or quantitative divergence could have significant impacts on postmating molecular interactions, only a few comparative studies have characterized the reproductive proteome composition and quantitatively compared individual proteins in populations/species with known PMPZ incompatibilities [21–23]. Furthermore, while PMPZ isolation is caused by incompatible interactions between the sexes, broad-scale efforts to identify interacting protein partners in conspecific crosses are rare [24], which impedes progress in predicting incompatible interactions in heterospecific crosses.

**Fig 1.**
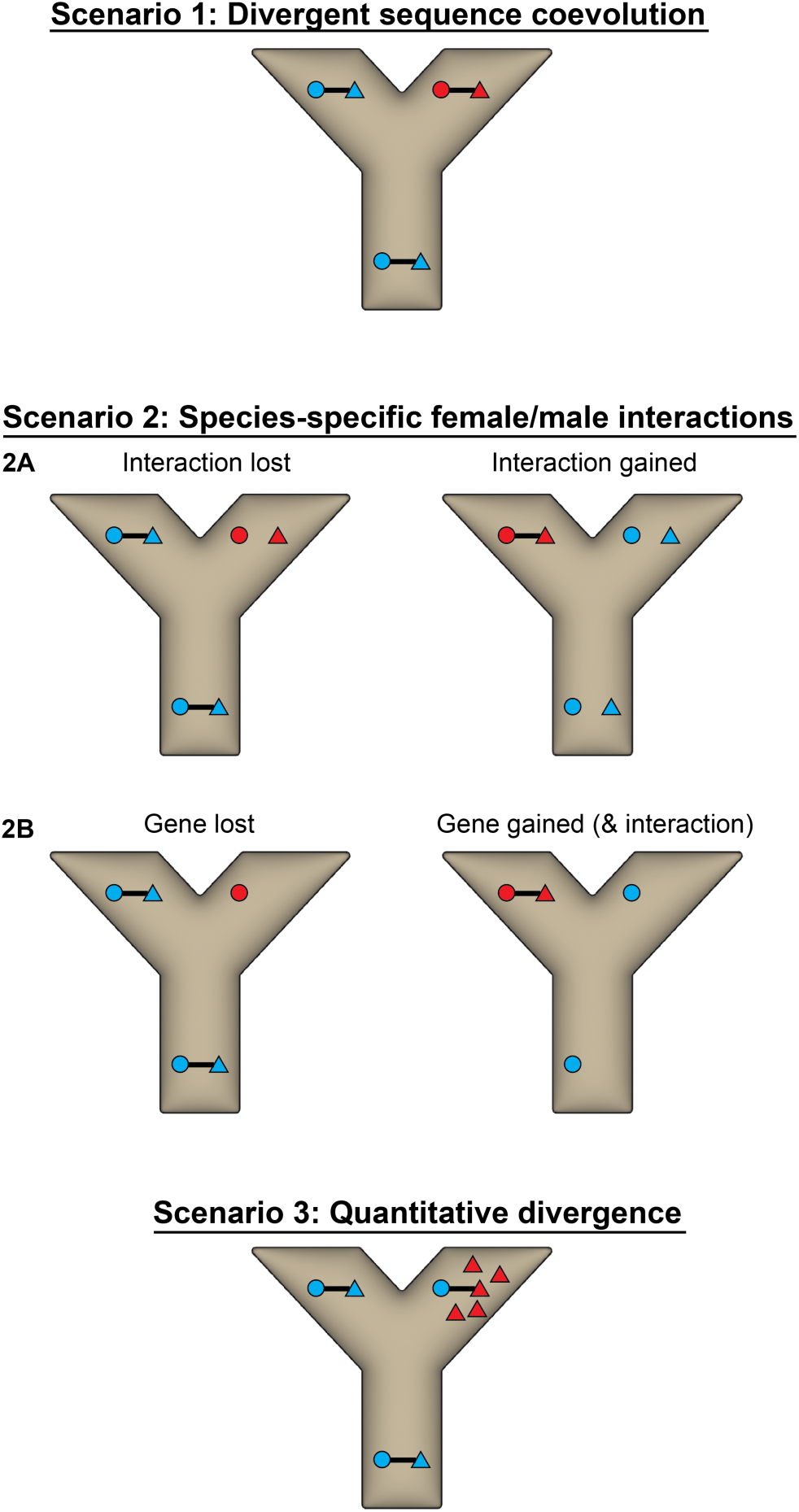
Divergent evolution of reproductive proteins in diverging lineages. Alternative scenarios describing how divergent evolution in lineages that split from a common ancestor could lead to PMPZ reproductive isolation. Circles and triangles represent different proteins. Blue indicates the ancestral condition, while red indicates the derived condition.

The *Drosophila mojavensis/arizonae* study system has a long history of speciation research [25]. Previous studies have shown that although crosses between these species exhibit several forms of reproductive isolation, PMPZ isolation is the only barrier that is consistently strong across all crosses that have been studied [25–28]. Females mated to heterospecifics exhibit sharp reductions in fertilization success and fail to efficiently degrade the insemination reaction mass, a clot-like mass that forms in the female reproductive tract during copulation [26,27]. Although the mass degrades over the course of several hours in females mated to conspecifics, it lasts for much longer in heterospecifically-mated females, and in some cases appears to persist indefinitely [26]. While the function of the insemination reaction has not been elucidated, it is hypothesized to mediate sexual conflict over female mating rate or be involved in facilitating the incorporation of male ejaculate components into female somatic tissues, as occurs in both species [29–31]. Previous research using expressed sequence tags identified candidate female reproductive tract proteins in *D. arizonae* [32], and the accessory gland proteome of *D. mojavensis* was characterized to predict candidate SFPs [33] Additionally, multiple studies have revealed widespread disruption of normal transcriptomic responses in heterospecifically-mated *D. mojavensis* and *D. arizonae* females [27,34,35] However, divergence in reproductive proteomes between the species, and the potential consequences for PMPZ isolation, has not been analyzed.

In this study, we adapt stable isotope labeling by amino acids in cell culture (SILAC) for use in whole flies to compare male and female reproductive proteomes in *D. mojavensis* and *D. arizonae* [36–40]. Specifically, our SILAC metabolic labeling strategy allows for unambiguous identification and robust quantitative comparisons of male SFPs. We further provide evidence for functional significance of three shared SFPs previously shown to be under divergent selection in *D. mojavensis* and *D. arizonae* [41]. Our design also allows semi-quantitative comparisons of female reproductive proteomes. Using these data, we leverage advances in predicting protein-protein interactions [42] to identify potential interactions between male and female proteases and protease inhibitors. We further predict disrupted interactions in heterospecific crosses and evaluate the relative contribution of divergence in protein sequence, proteome composition, and quantitative changes in contributing to incompatibilities. Overall, our results provide significant insights into the molecular basis of PMPZ reproductive incompatibilities.

## Results

### Extensive compositional, quantitative, and sequence divergence in male seminal fluid proteomes

The composition of seminal fluid proteomes diverged considerably between *D. mojavensis* and *D. arizonae*, with 26% of proteins identified being unique to one species (Fig 2A). Of these unique proteins, 93% had orthologs in the other species, indicating that recruitment/loss of proteins to seminal fluid is relatively common. Quantitative comparisons between 158 proteins with shared peptides also revealed extensive divergence, with 63% differing quantitatively between the species (Fig 2C). We also compared protein sequence divergence among shared SFPs, unique *D. mojavensis* SFPs, unique *D. mojavensis* SFPs, and genome-wide background. This analysis revealed that unique *D. mojavensis* and *D. arizonae* SFPs had higher Ka/Ks values than the genome background, while values for shared SFPs were intermediate and not significantly different from the unique SFPs or genome-wide estimates (Fig 2B). We also conducted phylogenetic tests of molecular evolution for all SFPs including the four *D. mojavensis* subspecies, genetically distinct northern and southern *D. arizonae* populations, and the closest relative to the *mojavensis/arizonae* cluster, *D. navojoa*. For aBSREL, we identified genes in which positive selection was inferred on branches separating all of *D. mojavensis* and *D. arizonae*. Positive site tests (FUBAR, BUSTED, and MEME) identified sites that were under diversifying selection across the phylogeny. Overall, 46% of SFPs were found to be under positive selection by at least one of these tests (Fig 2A; File S1).

**Fig 2.**
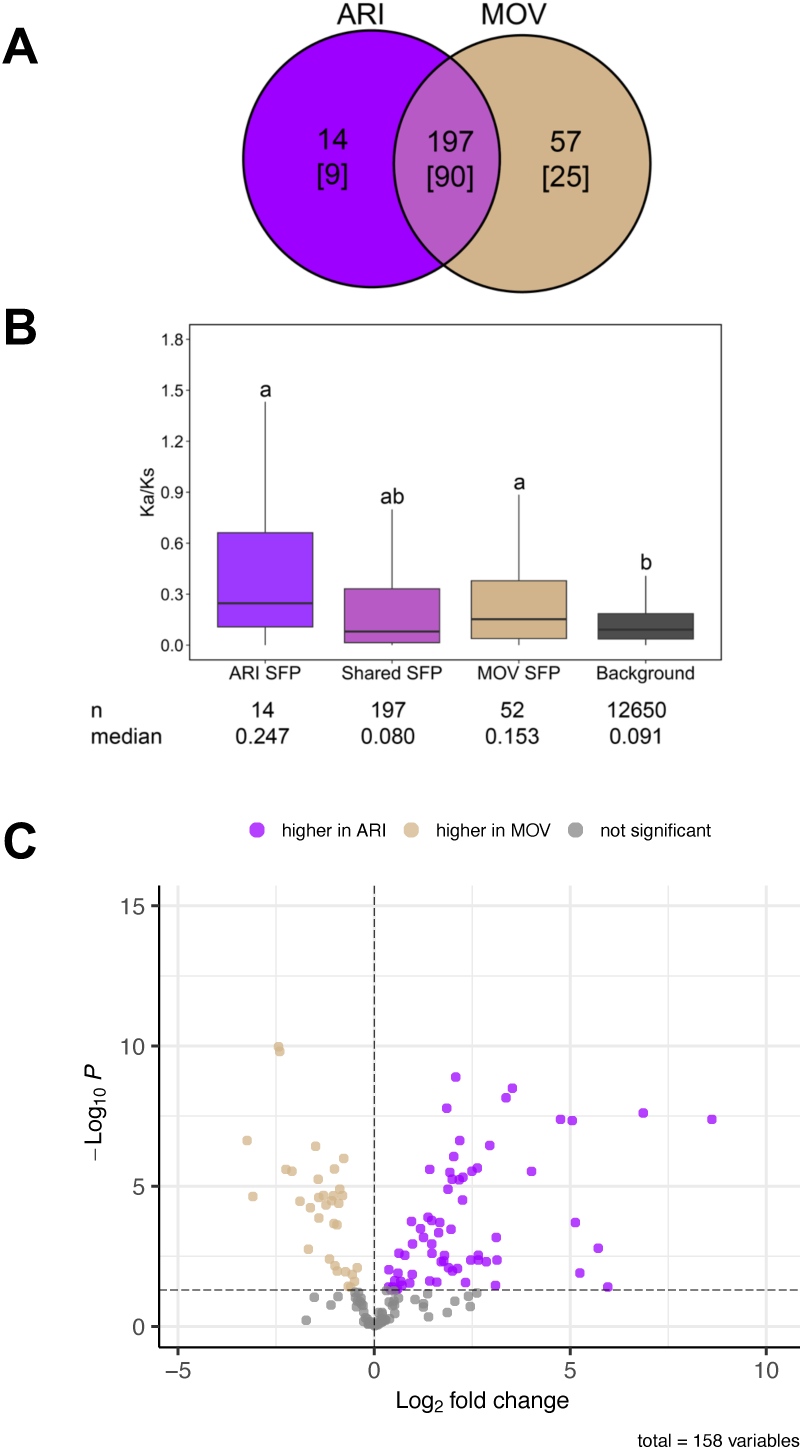
Compositional and quantitative divergence in seminal fluid proteomes. (A) Venn diagram showing shared and unique SFPs in *D. mojavensis* and *D. arizonae*. Numbers in brackets indicate the number genes with at least one HyPhy test suggesting positive selection. (B) Volcano plot showing quantitative differences between the species for individual SFPs. Only SFPs with shared peptides were included in the analysis. Dashed line represents significance threshold using FDR <0.05.

Enrichment analysis of SFP protein domains revealed mostly overlapping enriched categories between the species, although the number of genes in these clusters differed for some domains (Fig S1). Differences in enrichment of fibrinogens and papain-like cysteine peptidases are especially notable because a previous study showed that genes in these clusters affect the size of the insemination reaction mass and/or fertilization efficiency in *D. arizonae* [40]. Enriched domains also included protease inhibitors (e.g. serpins and Kunitz inhibitors) and CAP/CRISP proteins, which are known to play critical roles in reproductive processes in many species[43,44]. Furthermore, 50% of shared genes within enriched clusters differed quantitatively between the species (Fig 3) suggesting that quantitative differences may have functional consequences.

**Fig 3.**
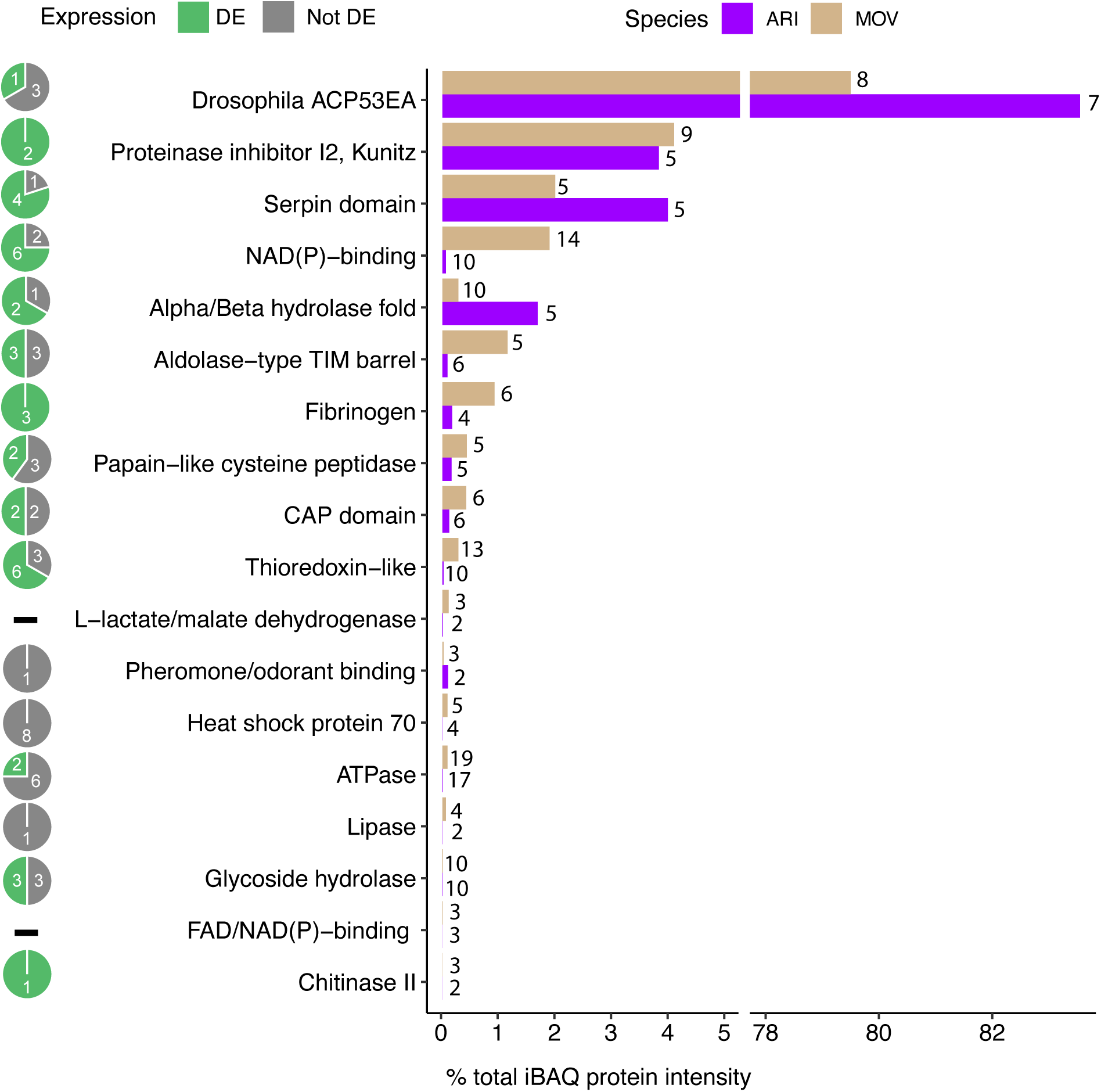
Quantitative analysis of enriched SFP protein domains. Quantitative comparison of enriched SFP protein domains in *D. mojavensis* and *D. arizonae*. Abundance of each domain was estimated by pooling iBAQ intensities for all proteins with a specific domain. Numbers above bars represent the total number of proteins in in the cluster. Pie charts show the proportion of individual genes in each cluster that were differentially expressed. This includes only genes in each cluster that had shared peptides between the species.

To better understand the relative quantitative composition of the seminal fluid proteomes, we used iBAQ quantification, which normalizes protein intensities based on the number of theoretical tryptic peptides, to compare the amounts of different enriched domains present within samples relative to total protein intensity. While not as robust as quantitative comparisons of single genes, this method gives a reasonable approximation of the relative abundances of different protein types within a sample. This analysis revealed the surprising finding that 80-84% of the male seminal fluid proteome was comprised of proteins with an ACP53EA domain (Fig 3). In *D. mojavensis*, MOV18622 comprised 57% of the total ACP53EA intensity, while MOV26980 made up an additional 17%. The rest was spread out among other paralogs with 5% being the highest total. The pattern was similar in *D. arizonae*, with ARI18622 and/or ACP2 (there were not enough unique peptides to distinguish them) comprising 56% of the total and ARI26980 making up an additional 24%. Protein-protein interaction predictions with Alphafold 3 suggest that the two most abundant paralogs in both species (18622 and 26980) likely form heterodimers (iptm=0.79 in *D. mojavensis* and 0.78 in *D. arizonae*), while neither paralog is predicted to form homodimers (all iptm values < 0.2). Of the remaining enriched domains, serine protease inhibitors, including serpins and Kunitz inhibitors, made up 6-8% of the total protein intensity, while all other domains constituted a relatively low proportion of the overall total (Fig 3). The diversity and quantity of serine protease inhibitors in male seminal fluid is interesting because we identified only four serine proteases in the male seminal fluid proteome.

### Most SFPs are also associated with sperm

To compare the composition of the seminal fluid proteome with the sperm-associated proteome, we performed LC-MS/MS on sperm samples isolated from the male seminal vesicle in each species. The majority of SFPs in both species (78% in *D. mojavensis*; 73% in *D. arizonae)* were also found in sperm samples (Fig S2). Although we cannot completely rule out that some male proteins in female reproductive tract samples came from sperm, our protein extraction method enriched for soluble, non-cellular proteins, so we believe this is an unlikely explanation. Unique SFPs in both species exhibited higher rates of molecular evolution compared to proteins shared in seminal fluid and sperm, sperm alone, and the genome-wide average (Fig S2).

### Divergently evolving SFPs are linked to postmating phenotypes in females

In a previous study, we identified four predicted SFPs that experienced divergent positive selection between *D. mojavensis* and *D. arizonae* (11629, 20219, 19546, and 23009) making them candidates for involvement in reproductive incompatibilities between the species[41]. One of these genes (*ARI20219*) is orthologous to *antares* in *D. melanogaster*, which functions in the sex peptide network and has demonstrated effects on female fertility[24] Transcripts of these genes are also transferred from males to females during mating, and our previous study found evidence for translation by females for three of the genes (i.e. male-derived female-translated proteins or mdFTPs) [34,40]. We further demonstrated that ARI11629 has functional effects on fertilization and the size of the insemination reaction mass in *D. arizonae* [40]. Our findings here confirm that these proteins are components of the seminal fluid proteome in both species, and molecular evolutionary analyses corroborate earlier findings indicating positive selection on 20219, 19546, and 23009 (11629 was excluded because it is not found in the genome of all seven populations/species used in our analyses).19546 was found in higher abundance in *D. arizonae,* while 23009 did not differ quantitatively between species (20219 did not have shared peptides).

To test for functional effects of these proteins on reproduction we used CRISPR to generate knockout (KO) lines for ARI20219, ARI19546, and ARI23009 in *D. arizonae*. We used these lines to test the effects of male KO mutations on female oviposition, larval hatching, and the formation/degradation of the insemination reaction mass. The results of these experiments indicated that KO mutations for all genes reduced larval hatching (Fig 4A). Furthermore, knockout of ARI20219 and ARI23009 in males resulted in a smaller insemination reaction in females (Fig 4B and 4C). Knockouts did not affect female oviposition for any gene (Fig 4D).

**Fig 4.**
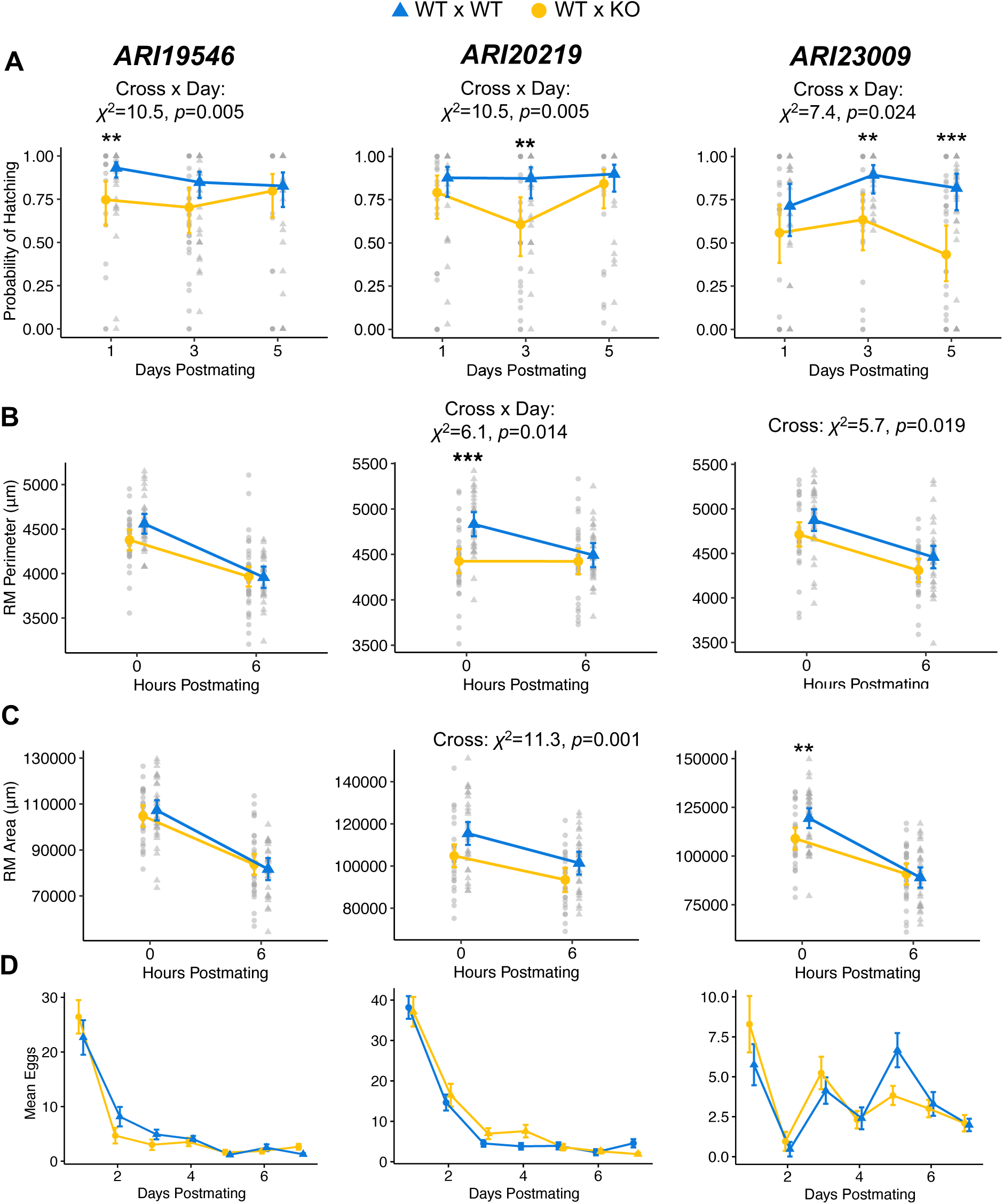
Knockout of male SFPs affects reproductive outcomes. Effect of male KO on hatching, oviposition, and insemination reaction mass for (A) *ARI19456*, (B) *ARI20219*, and (C) *ARI23009*. Data plotted in the background represent mean per vial in (A), and individual data points in (B) and (C). Significant interactions or cross effects are listed. If the interaction was significant, stars represent significant differences of estimated marginal means using Tukey’s method. \*\**p*< 0.01; \*\*\**p*<0.001.

### Female reproductive tract proteomes

Although we focused on identifying and quantifying male SFPs, our experimental design also enabled identification of unlabeled female reproductive tract proteins. Species-specific identifications are complicated by the fact that reproductive tracts of both species were combined before analysis, so we can only differentiate proteins with species-specific peptides. Considering all unlabeled proteins potentially produced by one or both species, we identified 1066 female reproductive tract proteins. Notably, 67% of SFPs were found to also be produced by females (Fig 5A). Overall, female proteins had lower rates of molecular evolution than unique male proteins (Fig 5B). Protein domain enrichment analysis identified 52 enriched domains (File S2). Most domains that were enriched in the seminal fluid proteome were also enriched in females, except for ACP53EA, L-lactate/malate dehydrogenase, CAP, fibrinogen, and chitinase II.

**Fig 5.**
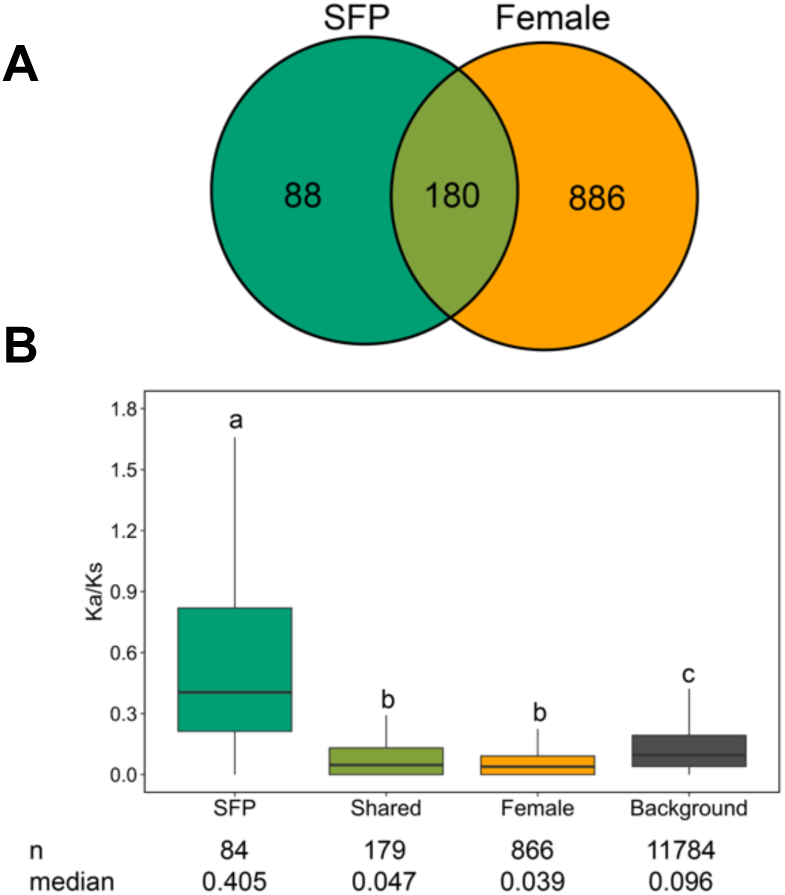
Comparison of composition and divergence of female and male reproductive proteomes. (A) Venn diagram showing overlap of male and female reproductive proteomes in *D. mojavensis* and *D. arizonae*. Female proteomes from both species are combined because female proteomes were not isotopically distinguishable. (B) Male specific SFPs exhibited higher Ka/Ks values than those shared with females or female only proteins, which evolved at lower rates than the genome-wide background. Groups under different letters were significantly different using *p*< 0.05.

While serine proteases were not enriched in the female reproductive tract, several were among the most abundant proteins, altogether making up ∼17% of the total female protein intensity. Since some serine proteases had species-specific peptides, we used these to identify and informally compare relative iBAQ quantities between the species. The most abundant proteases belonged to a cluster of paralogs we refer to as “SP cluster 1” (Fig 6). This cluster includes five paralogs in *D. arizonae* (ARI17775, ARI01606, ARI01588, ARI01589, and ARI01687), six in *D. mojavensis* (MOV17775, MOV02811, MOV02795, MOV02867, MOV23804, MOV02794). Since orthology was ambiguous, we compared relative quantities at the level of the cluster rather than individual proteins. Notably, outside of this cluster the next most abundant serine protease in *D. arizonae* (ARI01617) lacks an ortholog in *D. mojavensis* (Fig 6). This gene belongs to a group of paralogs that includes other highly abundant proteins including 26762, 26162, 00758, and 26169. Overall, *D. arizonae* females produced ∼3X the amount of serine proteases relative to *D. mojavensis*.

**Fig 6.**
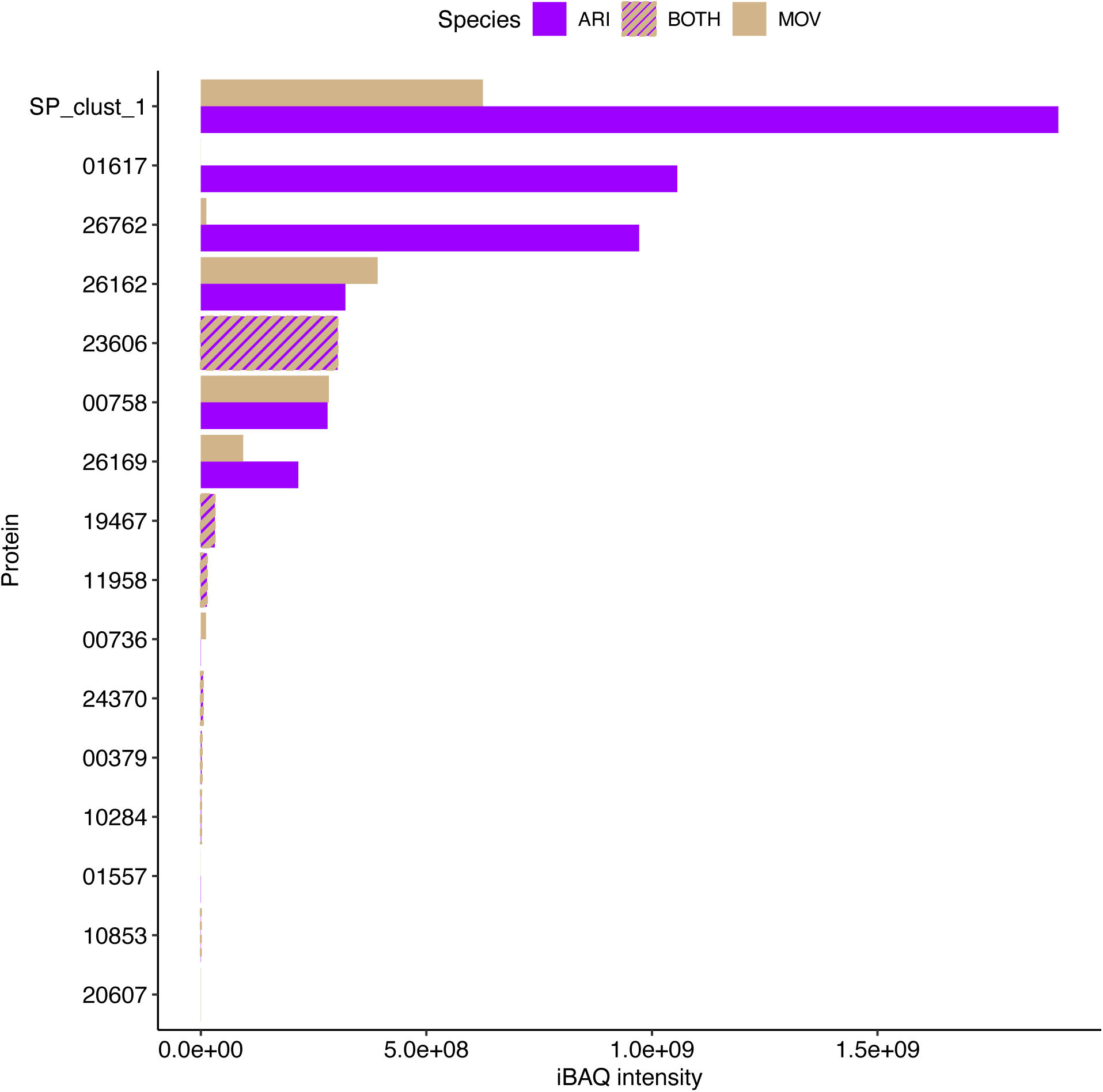
Extensive quantitative divergence in female serine proteases. Quantitative comparisons of female serine proteases in *D. mojavensis* and *D. arizonae* use iBAQ intensities of species-specific peptides. Hatched bars represent proteins that lacked species-specific peptides and thus could not be distinguished by species.

### Predictions of protein-protein interactions identifies male-female interacting partners

Given the high relative abundance and rapid evolution of serine proteases in females, together with the diversity and abundance of serine protease inhibitors in male seminal fluid, we further investigated potential interactions among proteases and inhibitors within and between the sexes. Specifically, we used AlphaFold 3 to model protein structures and predict interacting proteins within each species [42]. We included all male and female serine proteases, serpins, and Kunitz inhibitors. We also included cysteine peptidases, as these were enriched in female reproductive tracts and seminal fluid and can also be regulated by Kunitz inhibitors [45,46].

Using an iptm ≥ 0.80 to identify high confidence interactions, we found 265 predicted protein-protein interactions. This included 13 and 16 protease-protease interactions in *D. arizonae* and *D. mojavensis*, respectively, and 121 and 115 interactions between proteases and inhibitors in *D. arizonae* and *D. mojavensis*, respectively (File S3). Most protease-protease interactions were highly specific, with a single protease predicted to interact with 1-3 other proteases (Fig S3). While some interactions were conserved in both species, the overall network structure varied considerably (Fig S3). Interactions between serpins, which inhibit serine proteases, were also highly specific, with a single serpin predicted to interact with no more than two serine proteases (Fig S4). In contrast, Kunitz inhibitors displayed broad specificity, with most capable of inhibiting many serine proteases, and in some cases, cysteine peptidases (Fig S4). A total of 138 predicted interactions were between the sexes. This includes interactions between unique male and female proteins and cases where at least one of the interacting proteins is produced by both sexes.

### Predicted disruptions to protein-protein interactions in heterospecific crosses

To identify potentially disrupted interactions that might contribute to PMPZ isolation, we focused on the 138 intraspecific interactions between male and female proteins. We hypothesize that incompatible protein-protein interactions leading to PMPZ isolation may evolve under scenarios 1-3 presented in Fig 1. To evaluate evidence for scenario one, we modeled interspecies male-female interactions for conserved interactions present in both species. We considered interactions to be disrupted in heterospecific crosses when iptm ≤ 0.6. None of the 53 conserved interactions were predicted to be disrupted in either heterospecific cross. This supports a model of coevolutionary constraint rather than divergence (Fig 1; scenario 1), which is further bolstered by data from *D. navojoa* indicating that 85% (29/34) of these interactions were also conserved in this species at iptm ≥ 0.8 (19 interactions involved missing orthologs in *D. navojoa* and could not be evaluated) (File S3). This is true despite that fact that 72% of the genes involved in these interactions were predicted to be under positive selection by at least one comparative test. To evaluate evidence for scenario two, we modeled interspecific interactions for cases where predicted interactions were lineage specific (i.e. an interaction was gained or lost in one species). We considered interactions to be disrupted when (1) an interaction observed in conspecifics was lost when the male protein was heterospecific (disruptive loss), and (2) an interaction not observed in conspecifics was gained when the male protein was heterospecific (disruptive gain). We identified disruptions in 44% of these cases, including 12 disruptive losses and 18 disruptive gains (Fig 7). We also found evidence that male-female interactomes diverged through compositional changes to reproductive proteomes (Fig 1; scenario 2). Specifically, we identified 23 interactions involving four species-specific genes that fit scenario 2B. Considering the presence/absence of orthologs in *D. navojoa*, we inferred this resulted from two gene losses and two gene gains.

**Fig 7.**
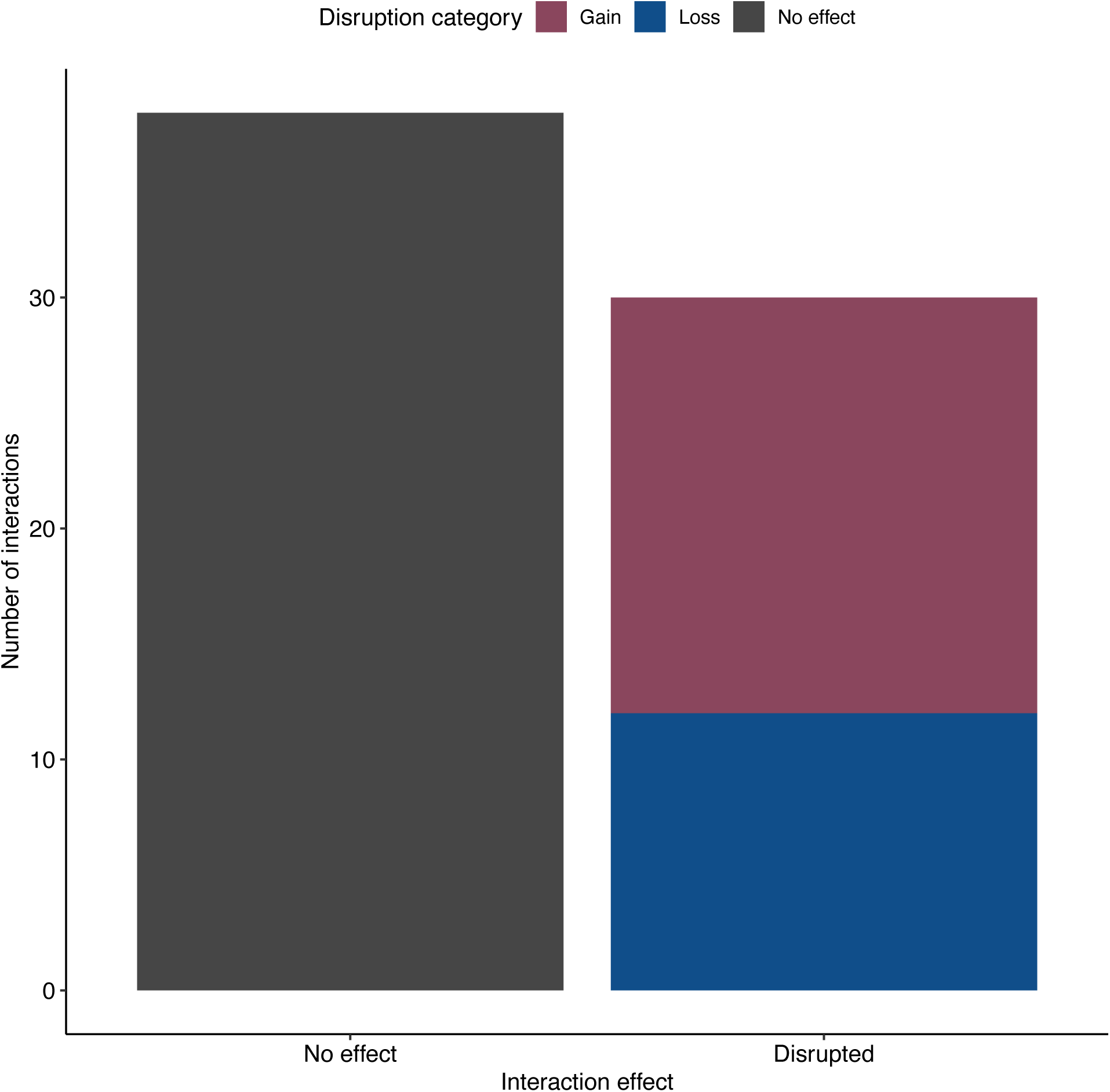
Compositional changes are predicted to result in disrupted male-female interactions in heterospecific crosses. The pattern of predicted disruption in heterospecific crosses for Fig 1, scenario 2. Disruptive losses represent cases where an interaction present in the intraspecific cross of one species was lost in the heterospecific cross with that female, while disruptive gains represent cases where an interaction that was not present in intraspecific cross of one species was present in the heterospecific cross with that female.

An additional 29 intraspecific interactions involved genes in paralogous clusters where gene orthology could not be determined. These interactions could therefore fit any of the scenarios described above. However, we do find some evidence for gains and/or losses (scenario 2), as five cases involve proteins that interact with proteins in the cluster in one species but not the other (one novel interactor is a lineage-specific gene).

## Discussion

Comparative analysis of *D. mojavensis* and *D. arizonae* seminal fluid proteomes revealed extensive divergence at the compositional, quantitative, and coding sequence levels. Approximately one quarter of SFPs were unique to one species, while 63% of conserved SFPs were present in different relative quantities, and almost half exhibited sequence evolution consistent with adaptive divergence. Furthermore, analysis of protein domains revealed divergence in proteins with known roles in driving the postmating response, suggesting observed changes are likely to have functional consequences for reproductive outcomes. For example, we previously found evidence for involvement of a fibrinogen-like protein (ARI26694) and a cysteine peptidase (ARI11629) in the formation and degradation of the insemination reaction mass and fertilization (ARI11629) in *D. arizonae*[40]. Both protein domains were enriched in the seminal fluid proteomes of *D. mojavensis* and *D. arizonae*, but exhibited lineage-specific compositional changes, quantitative changes, and divergent sequence evolution. Additional protein domains with established links to reproduction (CAP/CRISP and protease inhibitors) [43,44] also exhibited considerable divergence at all levels between the species.

Male gene knockout experiments with three divergent CAP/CRISP proteins provide direct evidence that these proteins play functional roles in reproduction in *D. arizonae*, further solidifying their potential involvement in PMPZ isolation. All three genes were also previously identified as candidate mdFTPs [40]. However, at this point, we cannot tease apart the effects of proteins transferred by the male vs. those produced by females from male RNA. Male KO of all these genes resulted in reduced fertilization success, which is consistent with known roles of CAP/CRISP proteins in fertilization in organisms ranging from insects to mammals [24,43]. Male KO of 2/3 genes also resulted in a smaller initial insemination reaction mass. While the mechanism by which they influence the size of the reaction mass is unclear, these proteins are related to Hymenopteran venom proteins that are known to induce allergic reactions [47]. Although the molecular origin of the insemination reaction mass is unknown, these results suggest the possibility that an inflammatory immune response could be involved.

Relative quantification of different seminal fluid proteins within species revealed the surprising finding that over 80% of the seminal fluid proteome for both species consisted of proteins with an ACP53EA domain. In *D. melanogaster*, ACP53EA is known to be involved in sperm storage and plays a functional role in sperm competition [48]. *ACP53EA* belongs to a multigene family found throughout *Drosophila,* although there is extensive copy number variation across the genus. *Drosophila mojavensis* and *D. arizonae* have eight copies in tandem on chromosome 5 (Muller element C). At least two of these are also candidate mdFTPs in *D. arizonae* [40]. Although all eight copies were SFPs in *D. mojavensis* and *D. arizonae*, 18622/ACP2 and 26980 constituted most of the protein intensity for the group. These small proteins have no conserved domains that are functionally characterized, but PPI modeling indicated that 18622/ACP2 and 26980 may form heterodimers. The significance of this is unclear, but the fact that these proteins are transferred in such high amounts suggests that the ability to form heterodimers could be important to their function.

The female reproductive proteome substantially overlapped the seminal fluid proteome, with 67% of SFPs also being produced by females. This finding is consistent with recent studies in *D. melanogaster* demonstrating that many SFPs are not male-specific, as was previously assumed, but rather exhibit shared expression in male and female reproductive tracts [49,50]. Expression of SFPs is particularly high in female sperm storage organs, likely reflecting the need for females to maintain sperm viability during storage [49]. We also observed higher rates of molecular evolution in male-specific SFPs, while shared SFPs and female-specific proteins had lower rates of molecular evolution than the genome background. These results also corroborate recent work in *D. melanogaster*, which has shown that subsets of SFPs experience different selection pressures, with male-specific proteins evolving the most rapidly [49]. Shared expression by both sexes may constrain the evolution of reproductive proteins.

The most abundant female reproductive proteins were serine proteases belonging to three small gene families expressed exclusively in *D. mojavensis/D. arizonae* female reproductive tracts. Kelleher and Pennington [51] and Kelleher et al. [32] demonstrated extensive divergent evolution of these in *D. mojavensis* and *D. arizonae*, including lineage-specific duplications, gene conversion, and adaptive sequence evolution. Serine proteases play important roles in reproduction in *Drosophila* and other organisms, including sperm storage and capacitation, immunity, mating plug formation/degradation, and influences on female behavior [44], which highlights the potential importance of these proteins to reproduction in *D. mojavensis* and *D. arizonae*.

We identified numerous predicted interactions among serine proteases, cysteine peptidases, and protease inhibitors, including intra- and intersexual interactions. We used predicted interactions among proteases, and among proteases and inhibitors, as a system to test hypotheses about predicted causes of reproductive incompatibilities (Fig 1). Surprisingly, we found no evidence to support a coevolutionary model where divergent sequence evolution of interacting proteins results in incompatible interactions in heterospecific crosses. Although 72% of proteins involved in conserved intersexual interactions appear to be under divergent selection in *D. mojavensis* and *D. arizonae*, none of these interactions were predicted to be disrupted in heterospecific crosses. These findings lend support to recent arguments claiming that much of the protein sequence divergence observed in reproductive proteins may result from non-adaptive processes rather than from positive selection driven by postcopulatory sexual selection [10–12]. Our data suggest that PMPZ incompatibilities may instead be most likely to arise from compositional changes to reproductive proteomes and/or interactomes that result from divergent gains/losses of genes or interactions. We note that this does not necessarily mean that sexual selection and sexual conflict are not important drivers of PMPZ incompatibilities, as changes to the interactome through gains and losses of genes and/or interactions could be driven by these processes.

These conclusions are subject to a few caveats. Most importantly, our conclusions rely on the accuracy of PPI interaction modeling rather than direct experimental evidence for interactions. AlphaFold 3 is generally highly accurate, especially when there is abundant structural information for proteins of interest (e.g. proteases and inhibitors), and we used conservative cut-offs to identify interactions [42]. Nonetheless, our results require future validation with experimental methods. We also note that while AlphaFold 3 predicts the structure and likelihood of proteins binding, it does not predict the strength and efficiency of the interaction. Thus, although heterospecific proteins may structurally interact, the strength or efficiency of the interaction may be compromised. Finally, our conclusions are based on results derived exclusively from proteases and inhibitors, which may not be representative of other types of interactions that are crucial for successful reproduction. For example, we demonstrated the involvement of three divergently evolving CAP/CRISP proteins in several postmating phenotypes. Unfortunately, we were unable to perform PPI modeling for this class of proteins due to a lack of *a priori* information about potential interacting partners. Ultimately, testing a broader set of interactions, *in silico* and/or *in vivo*, is necessary to draw more robust conclusions about the effects of divergent protein sequence evolution on protein-protein interactions. Despite these limitations, our results lay a solid foundation for future experimental research aimed at identifying intersexual molecular interactions that drive PMPZ reproductive incompatibilities.

Although we cannot directly test the extent to which quantitative differences reproductive proteomes are likely to cause incompatibilities (Fig. 1), the scope for such effects is large. Indeed, 63% of the male seminal fluid proteome differed quantitatively between species, and quantitative divergence in female proteins also appears to be extensive. Quantitative differences between species in proteases and their inhibitors may be especially likely to contribute to reproductive incompatibilities due to misregulation of protease activity, which is extensive [44], in heterospecific crosses. *Drosophila mojavensis* and *D. arizonae* differed substantially in protease production, with differences being especially pronounced for female serine proteases (Fig. 6). We also observed differences in the overall level of male inhibitors (Fig. 3).

Furthermore, all male inhibitors included in the quantitative analysis of male seminal fluid (four serpins and two Kunitz inhibitors) were differentially expressed. Both Kunitz inhibitors and 3/4 serpins were expressed at higher levels in *D. arizonae*, with the Kunitz inhibitors having the largest fold changes among all differentially expressed proteins (01590 fc= 388; 17759 fc= 111). Proper regulation of proteolysis is achieved by balancing the stoichiometry of proteases and inhibitors, and dysregulation of these processes is implicated in numerous pathologies related to many fundamental physiological processes [52–55]. The extensive quantitative divergence we found here may thus drive the evolution of reproductive incompatibilities, especially given the known importance of proteolysis regulation specifically in reproduction [44].

## Conclusions

Overall, our results demonstrate extensive divergence in reproductive proteomes at protein coding sequence, proteome/interactome composition, and quantitative levels. Although previous research has largely centered on protein-coding sequence divergence as the primary driver of PMPZ incompatibilities, our findings reveal that a broader perspective—encompassing both compositional and quantitative divergence—is essential for understanding the complex molecular basis of PMPZ incompatibilities. Altogether, these findings provide significant insights into the genetic evolution of PMPZ isolation, which is increasingly recognized as an important early-evolving reproductive isolating barrier.

## Methods

### Fly metabolic labeling

We used genome sequenced stocks of *D. mojavensis* (National *Drosophila* Species Stock Center (NDSSC): 15081-1353.01) and *D. arizonae* (NDSSC: 15081-1271.41) for all proteomic analyses. To differentiate male and female proteins we used a SILAC labeling strategy, which is generally considered the most robust method for quantitative proteomic comparisons [56]. We labeled *D. mojavensis* males with L-lysine-^13^C ^15^N (Lys8; Cambridge Isotopes Laboratory, Inc, Tewksbury, MA), *D. arizonae* males were labeled with L-lysine 4,4,5,5-D_4_ (Lys4; Cambridge Isotopes Laboratory, Inc, Tewksbury, MA), and females were unlabeled. Mated female reproductive tracts of both species were combined prior to proteomic analysis. The advantage of this approach is that it allows *D. mojavensis* and *D. arizonae* male proteins to be unambiguously identified within the female reproductive tract and compared quantitatively.

Although female proteins can also be distinguished from male proteins, species-specific designations are more challenging since unlabeled female tracts of both species had to be combined prior to analysis. As originally described in [40], we first reared a lysine auxotrophic strain of *Saccharomyces cerevisiae* (MATalpha leu2Δ0 lys2Δ0 ura3Δ0 in background BY4729; Dharmacon, Inc. Lafayette, CO) in synthetic liquid media consisting of yeast nitrogen base without amino acids, yeast synthetic lysine drop-out medium, and isotopically labeled lysine (Lys4 or Lys8). Cultures were incubated with shaking for ∼24 hours at 30°C. We pelleted cells by centrifugation and washed with sterilized water and centrifuged again. Pellets were lyophilized, and frozen at -20°C until used in experiments.

We prepared food as described in [40] by combining Lys8 or Lys4 labeled yeast, yeast nitrogen base without amino acids, pure molasses, agar, sterile water and methylparaben dissolved in ethanol. The food was autoclaved before dispensing into sterile glass vials in a biosafety cabinet. To collect eggs, adults of both species were placed in population cages overnight. Eggs were collected the next day and sterilized/dechorionated by soaking in 2.5% sodium hypochlorite for three minutes before placing in food vials with isotopically labeled yeast (∼100 eggs/vial). Newly emerging males were collected within a few hours of eclosion and were held on food consisting of pure molasses, agar, water, and a flake of lyophilized Lys8 or Lys4 labeled yeast. Flies were transferred daily to new vials for eight days to ensure they had reached reproductive maturity. This rearing protocol produced high labeling efficiency (99%) [40]. *Drosophila mojavensis* and *D. arizonae* females were collected within a few hours of emergence from standard banana-molasses food [57] with no isotopically labeled yeast.

The morning of experiments, labeled males of each species were combined with respective unlabeled females and copulations were observed. Mated females were placed in liquid nitrogen immediately after mating and were held at -80°C. Female reproductive tracts were removed in groups of five and placed in 20 μl ice cold 50mM ammonium bicarbonate. To enrich for extracellular secreted proteins, tracts were centrifuged at 17,000 rcf, and the supernatant was collected and stored at -80°C for analysis. We collected six biological replicates for each species.

### Sperm sample preparation

Sperm were collected from ∼20 unmated sexually mature flies of each species by tearing open the seminal vesicles and removing sperm clumps. Sperm were rinsed twice with Ringer’s solution and pelleted by centrifugation at 17,000 rcf. Pellets were immediately frozen in liquid nitrogen and stored at -80°C.

### Liquid chromatography tandem mass spectrometry (LC-MS/MS)

LC-MS/MS was performed at the Central Analytical Mass Spectrometry Facility and W.M. Keck Foundation Proteomics Resource at University of Colorado Boulder. Mated lower female reproductive tracts of *D. mojavensis* and *D. arizonae* were denatured, reduced and alkylated with 5% (w/v) sodium dodecyl sulfate (SDS), 10mM tris (2-carboxyethyl) phosphine hydrochloride (TCEP-HCl), 40mM 2-chloroacetamide, 50mM Tris pH 8.5 and boiled at 95°C for 10 minutes. The SP3 method[58] was utilized to prepare samples for mass spectrometry analyses. Protein lysates were treated with carboxylate-functionalized speedbeads (Cytiva) and acetonitrile was added to 80% (v/v) to promote binding to the beads. Beads were then washed twice with 80% (v/v) ethanol and twice with 100% acetonitrile. Digestion was performed with LysC/Trypsin mix (Promega) at a 1:50 protease to protein ratio in 50mM Tris pH 8.5. Samples were incubated and rotated at 37°C overnight. Digests were purified with an Oasis HLB 1cc (10mg) cartridge (Waters) following the manufacturers protocol and then dried using a vacuum rotatory evaporator. Peptides were directly injected onto a Waters M-class column (1.7 μm, 120A, rpC18, 75 μm x 250mm) and eluted from 2% to 20% acetonitrile with 0.1%(v/v) formic acid for 100 minutes then 20% to 32% acetonitrile for 20 minutes at 0.3 μl/minute using a Thermo Ultimate 3000 UPLC (Thermo Scientific). Mass spectrometry was performed with a Thermo Q-Exactive HF-X mass spectrometer (Thermo Scientific). MS1 scans were performed at 120,000 resolution from 380 to 1580 m/z with a 45ms fill time and 3E6 AGC target. For MS2 scans, the 12 most intense peaks were isolated with 1.4 m/z window with 100ms fill time and 1E5 AGC target and 27% HCD collision energy at 15,000 resolution. Dynamic exclusion was enabled for 25 seconds.

### Identification and quantitative comparison of SFPs

Proteins were identified and quantified using MaxQuant[59], assuming a triple SILAC labeling experiment (heavy [Lys8] =*D. mojavensis* SFPs, medium [Lys4] =*D. arizonae* SFPs, light [Lys] =*D. mojavensis* and *D. arizonae* female proteins). The protein database comprised combined *D. mojavensis* (NDSSC: 15081-1353.01) and *D. arizonae* (NDSSC: 15081-1271.41) proteins [60]using the longest transcript for each species (cactusflybase R2024_09). The analysis included the possibility of common variable modifications (Oxidation (M); Acetyl (Protein N-term) and PSM and protein FDR were set to 0.01. We required at least two peptides for a protein to be identified. Because variation in ionization efficiencies of different peptides can reduce the reliability of abundance estimates, we filtered our SFP dataset for quantitative comparisons to include only peptides shared between the species. This enabled us to quantitatively compare 158 of 271 total SFPs. Comparisons were made using the R package ‘artMS’ [61], which relies on the ‘MSstats’ v4.0 package [62] for statistical comparisons. We used the ‘equalize medians’ method of normalization and did not impute missing data. All mass spectrometry proteomics data have been deposited on the ProteomeXchange Consortium via the PRIDE [63] partner repository with the dataset identifier PXD065290 (temporary reviewer credentials, Username; Password: hCWEBXXmMQtc). All MaxQuant analysis parameter, database and output files are available at OSF (https://osf.io/e9waj/?view_only=e52d5b6dff874b0ab0c63805a5ababed).

### Molecular evolution

We downloaded protein sequences from all species from previously sequenced genomes (R1.0) [60] available on cactusflybase R2024_09. AGAT v1.0 [64] was used to obtain the longest transcripts for each gene and PRANK [65] was utilized to generate codon-based alignments of orthologs. Two sets of alignments were generated, a *D. mojavensis* (NDSSC: 15081-1353.01) and *D. arizonae* (NDSSC: 15081-1271.41) orthologs set (for the Ka/Ks calculation) and a set comprising sequences from seven cactophilic *Drosophila* including *D. mojavensis* (four populations), *D. arizonae* (two populations) and *D. navojoa* [60]. Estimates of Ka/Ks were performed using KaKs_Calculator v2.0 [66]. Tests for significant differences in Ka/Ks between groups were performed using pairwise Kruskal-Wallis tests in R. We used the HyPhy v2.5.60 [67] platform to execute tests of molecular evolution, aBSRE L[68] , BUSTED [69], MEME [70] and FUBAR [71]. For the latter three tests the alignments were partitioned by recombination blocks using GARD [72]. The entire phylogeny was designated as the foreground branch for BUSTED. Phylogenetic relationships determined by Benowitz et al. [60] were used for all these tests.

### Protein domain enrichment analysis

Identification of protein domains for *D. mojavensis* (NDSSC: 15081-1353.01) and *D. arizonae* (NDSSC: 15081-1271.41) was individually generated using InterProScan v5.69-101.0 [73]. Determination of enrichment of protein domains, based on IPR, was performed using the R package ‘WebGestaltR’ [74] with the default analysis parameters except setting the minimum group size to 8. A species-specific genome-wide background IPR dataset was used for the analysis of each species.

### Knockout experiments

Since rapid protein sequence evolution is hypothesized to drive the evolution of PMPZ incompatibilities (Fig 1; scenario 1), we sought to test the reproductive function of three shared SFPs (20219, 23009, 19546) that we previously demonstrated were under divergent selection in *D. mojavensis* and *D. arizonae* and were also identified as male-derived, female-translated protiens (mdFTPs) [40]. We used CRISPR knockouts in male *D. arizonae* (NDSSC: 15081-1271.41) to test the effects of these proteins on female fecundity, larval hatching, and the formation/degradation of the insemination reaction mass.

We designed two sgRNAs targeting the 5’ end of each gene following Bassett et al.[75] Embryos were injected into *D. arizonae* (NDSSC: 15081-1271.41) embryos by Rainbow Transgenic Flies, Inc. (Camarillo, CA). We established two different homozygous KO lines for each gene (*ARI19546*: 11bp and 38bp deletion; *ARI20219*: 5bp and 2bp deletion; *ARI23009*: 5bp and 13 bp deletion), which were crossed to generate transheterozygous males (to control for off-target effects) for use in experiments.

We paired unmated wild-type (WT) or KO males with WT females from a different *D. arizonae* line (ARTU2) and copulations were observed. For the fecundity assay, mated females were separated from males and moved to new food every 24 hours. Eggs were counted for seven days postmating. For the hatching assay, mated females were placed singly (*ARI20219* and *ARI23009*) or in groups of three (*ARI19546*) and rotated to new vials every 24 hours. We counted eggs on days one, three, and five, and counted the number of hatching larvae on subsequent days. Although this analysis does not differentiate between unfertilized eggs and embryonic mortality, none of these genes are expressed in developing embryos (cactusflybase.arizona.edu), so effects on embryo viability are unlikely. For the insemination reaction assay, we froze mated females in liquid nitrogen either immediately after mating or six hours postmating. We photographed lower reproductive using a camera mounted to a Leica S9i dissecting microscope (37.5X magnification). To measure the size of the insemination reaction we used the freeform drawing tool in ImageJ[76] to trace the outline of the lower female reproductive tract. We compared the perimeter and area, converting pixels to μm using photo resolution.

Fecundity data were analyzed as a negative binomial generalized linear mixed model with log link function using the R package ‘glmmTMB’ [77]. The factorial model included male genotype, day, and their interaction, and individual was considered a random effect. We used a binomial generalized linear mixed model with logit link function implemented in the R package ‘afex’[78] to analyze hatching data. The factorial model included male genotype, day, and their interaction, and vial was treated as a random effect. Anova tables for fecundity and hatching data were generated using the ‘car’ package [79] Insemination reaction data were analyzed as a factorial Anova using the ‘afex’[78] package, with a model that included male genotype, time, and their interaction as factors. *Post-hoc* comparisons from all experiments were analyzed using Tukey’s adjustment for multiple comparisons implemented in the ‘emmeans’[80] package in R.

### Quantitative comparisons of female serine proteases

Given previous studies suggesting that *D. mojavensis/D. arizonae* female serine proteases are important players in the female postmating response [32,51], we sought to quantitatively compare these proteins between the species. Since female proteins of both species were unlabeled and combined prior to analysis, we used species-specific peptides for comparisons. Specifically, we used iBAQ quantifications for these comparisons, which normalizes by dividing total protein intensity by the number of theoretically observable peptides [59,81]. While iBAQ quantification is intended to permit comparisons of the relative amounts of different proteins in a sample, we note the caveat that this method does not control for different peptide ionization efficiencies and is therefore likely to be less reliable than SILAC quantifications of common peptides (as used for male proteins). Nevertheless, we consider these estimates a reasonable approximation of the relative amounts of different female proteins in our samples.

### Protein-protein interaction modeling

We used AlphaFold 3 [42] to predict protein-protein interactions within and between the sexes in each species. We included all male and female serine proteases and inhibitors (serpins, Kunitz, and one Cathepsin I inhibitors). In addition, we also included cysteine peptidases as they were also enriched in male seminal fluid and female reproductive tracts and can be regulated by Kunitz inhibitors [45,46]. We first used Interpro searches to identify and remove signal sequences when annotations were available. Moreover, since many proteases are synthesized as zymogens, we included both active and inactive forms of the protease if cleavage sites were annotated. Following recommendations by the developers of AlphaFold 3 [42], we considered any models with Interphase Predicted Template Modeling (iptm) values ≥ 0.8 to be highly confident interactions.

We further investigated the subset of interactions predicted between male and female proteins for evidence of disruption in heterospecific crosses. Reciprocal interactions between the proteins of each species were modeled and interactions were considered disrupted if iptm ≤ 0.6. To evaluate the conservation of predicted interactions, we also modeled male-female interactions of predicted orthologs in *D. navojoa* using an iptm ≥ 0.8 cutoff.

## Acknowledgements

We thank Jeffrey Callan, Nathan Dee, Margaret Kirkpatrick, and Seth Nolen for assistance with establishing knockout mutation lines. We thank Christopher Ebmeier from the Central Analytical Mass Spectrometry Facility and W.M. Keck Foundation Proteomics Resource at University of Colorado Boulder for assistance with mass spectrometry. We also thank Juan Hurtado for providing comments on an earlier version of the manuscript.

## Supporting information

**Fig. S1.**
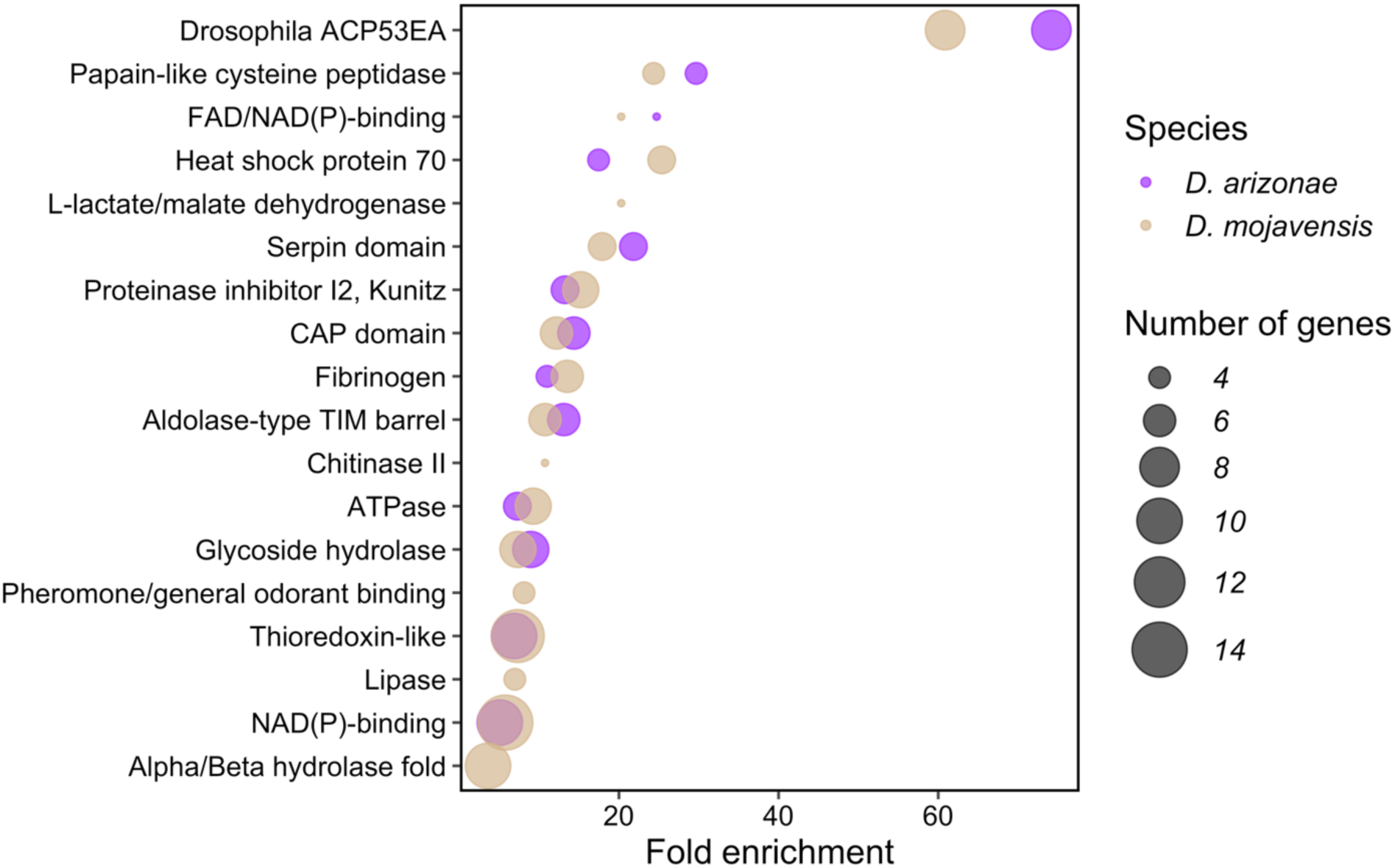
Enriched protein domains in seminal fluid proteomes. Enrichment of seminal fluid protein domains in *Drosophila mojavensis* and *D. arizonae* is largely overlapping, although there are differences in the number of genes associated with domains known to play a role in reproduction (e.g. fibrinogen and papain-like cysteine peptidase).

**Fig. S2.**
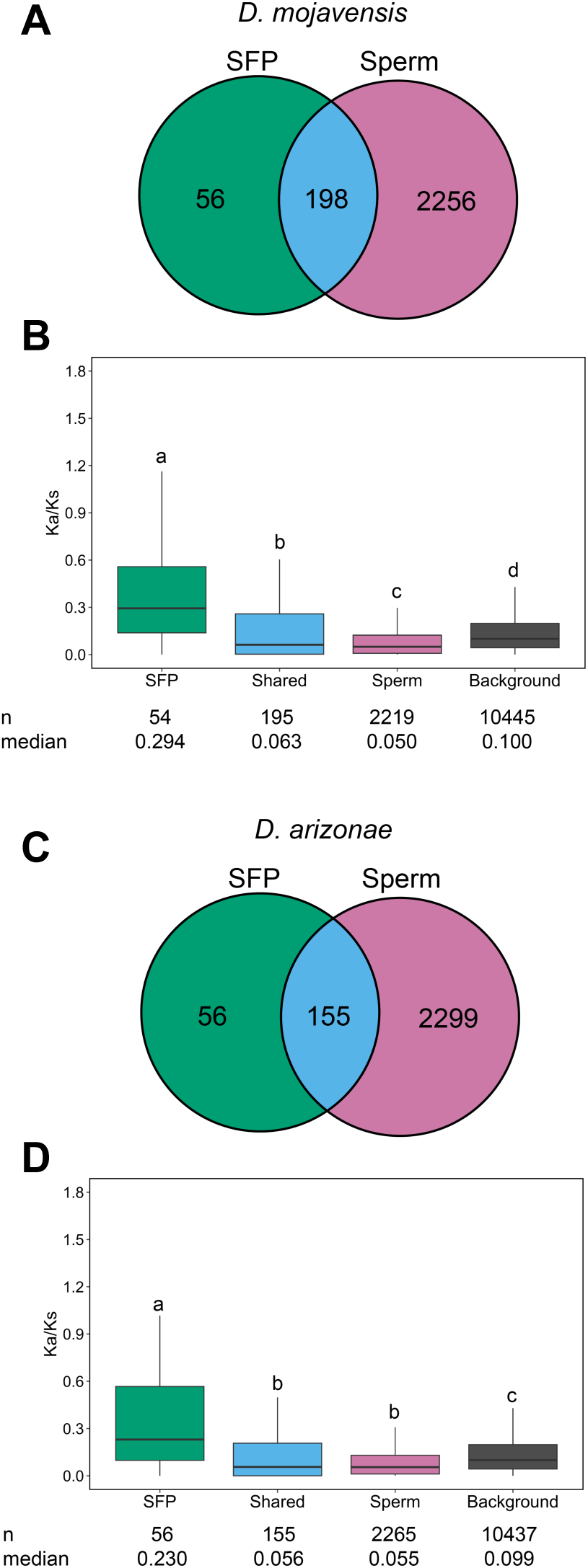
Most SFPs were also associated with sperm. (A) Venn diagram showing overlap between seminal and sperm-associated proteomes in *D. mojavensis*. (B) *D. mojavensis* SFPs unique to seminal fluid exhibited higher Ka/Ks values than those also found in sperm, sperm-specific, and genome background. Groups under different letters were significantly different using *p*< 0.05. (C) Venn diagram showing overlap between seminal and sperm-associated proteomes in *D. arizonae*. (D) *D. arizonae* SFPs unique to seminal fluid exhibited higher Ka/Ks values than those also found in sperm, sperm-specific, and genome background. Groups under different letters were significantly different using *p*< 0.05.

**Fig. S3.**
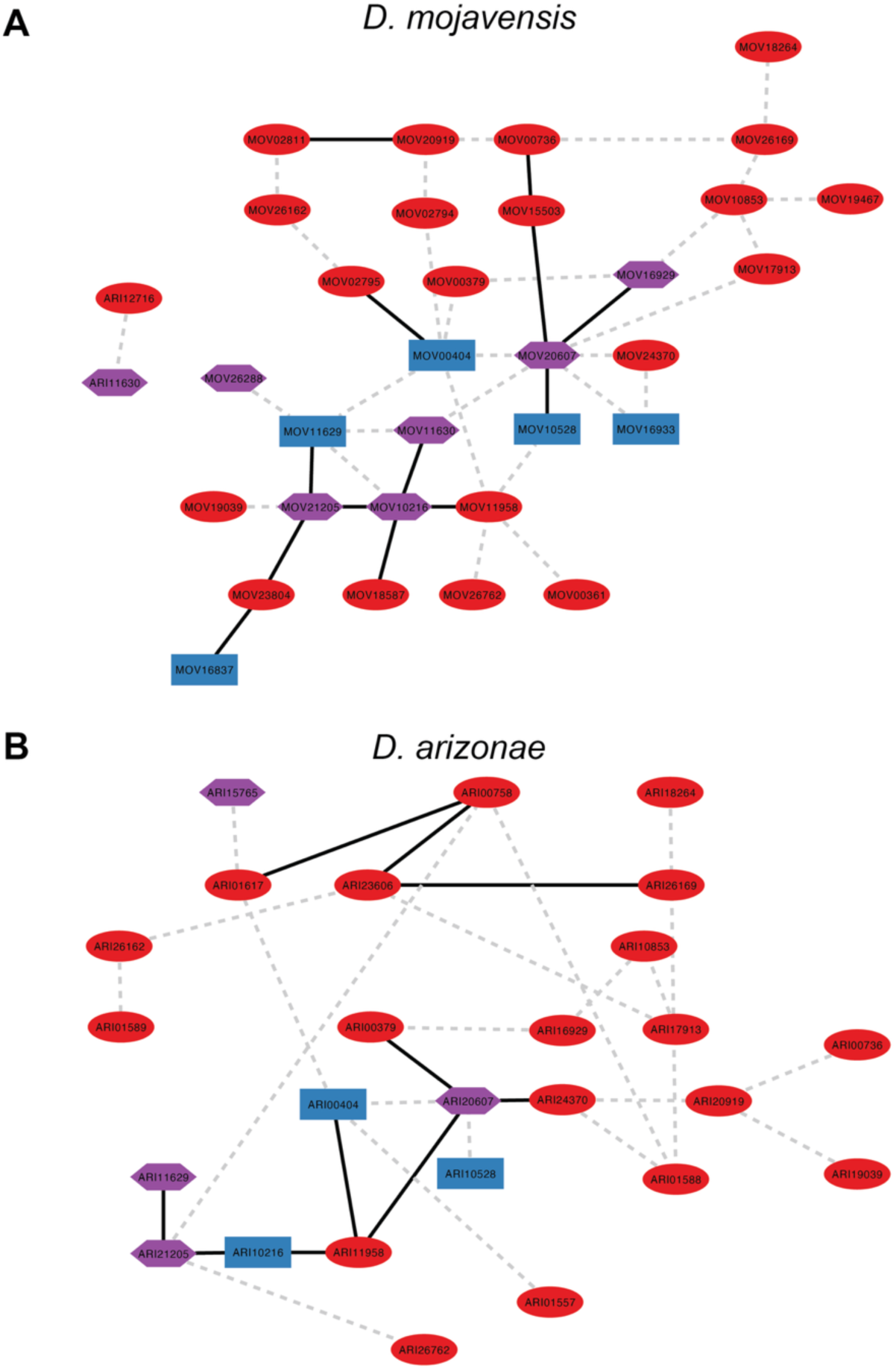
Divergence in protease interactions between *D. mojavensis* and *D. arizonae*. Protease interaction networks in (A) *D. mojavensis* and (B) *D. arizonae*. Solid lines indicate interactions with iptm≥ 0.8 while dashed lines indicate interactions with iptm≥ 0.7. Red circles= female-specific proteins; blue squares= male-specific proteins; purple hexagons= proteins produced by both sexes.

**Fig. S4.**
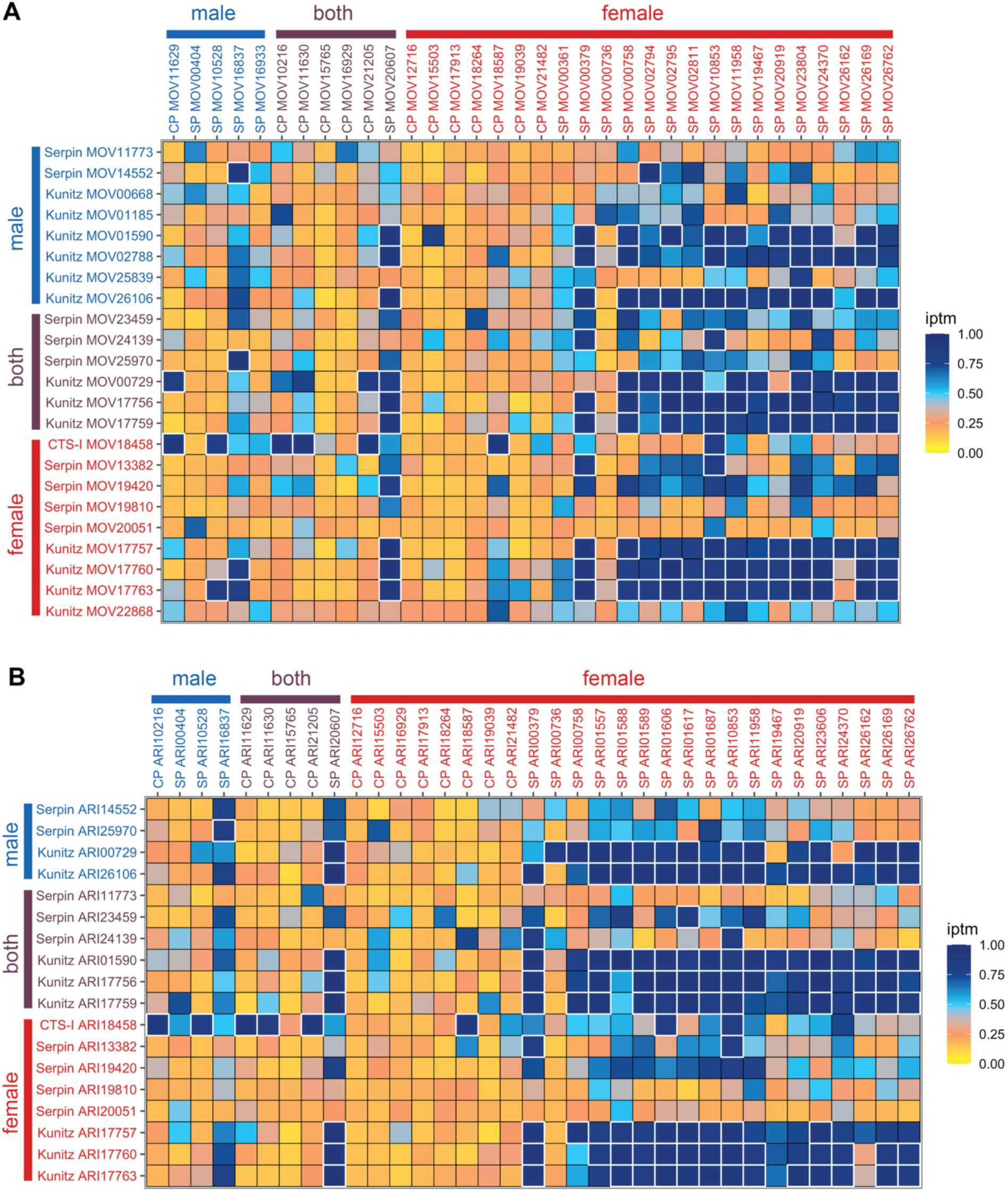
Predicted interactions between proteases and protease inhibitors. Heat map showing iptm values for interactions between proteases [Cysteine peptidases (CP) and Serine proteases (SP)] and protease inhibitors [Serpins, Kunitz and Cathepsin I (CTS-I)] for (A) *D. mojavensis* and (B) *D. arizonae*. Proteins with red labels are female-specific, proteins with blue labels are male-specific, and proteins with purple labels are produced by both sexes. Boxes outlined in white are highly supported interactions (iptm≥ 0.8).

**File S1. Comparative tests of molecular evolution.** Results for aBSREL, BUSTED, FUBAR, and MEME tests on male seminal fluid protein genes.

**File S2. Female protein domain enrichment**. List of enriched domains in *D. mojavensis/D. arizonae* female reproductive proteins.

**File S3. Results of protein-protein interaction analyses**. Results of analyses of protein-protein interactions using AlphaFold 3. Signal sequences were removed from protein sequences if annotated by InterPro. If annotated in InterPro, protein names that include “active” indicate the activated form of the protein.

